# Regional Disparities of Antenatal Care Utilization in Indonesia

**DOI:** 10.1101/793802

**Authors:** Agung Dwi Laksono, Rukmini Rukmini, Ratna Dwi Wulandari

## Abstract

**Introduction:** The main strategy for decreasing maternal morbidity and mortality with antenatal care (ANC). ANC aims to monitor and maintain the health and safety of the mother and fetus, detect all complications of pregnancy and take the necessary actions, respond to complaints, prepare for birth, and promote healthy living behavior. The study aims to analyze inter-regional disparities in ≥4 ANC visits during pregnancy in Indonesia.

**Methods:** Data sources from 2017 Indonesian Demographic and Health Survey (IDHS). With an analysis unit of women aged 15-49 years old, a sample of 15,351 women was obtained. Besides ANC as the dependent variable, other variables analyzed were place of residence, age, husband/partner, education, parity, wealth status, and health insurance. Analysis using Binary Logistic Regression for the final test to determine disparity.

**Results:** All regions show a gap with the Papua region as a reference, except the Maluku region which was not significant shows differences in the use of ANC compared to the Papua. Women in the Nusa Tenggara have 4,365 chances of making ≥4 ANC visits compared to the Papua region. Women in Java-Bali have 3,607 times more chances to make ≥4 ANC visits than women in the Papua region. Women in Sumatra have 1,370 chances of making ≥4 ANC visits compared to women in the Papua region. Women in Kalimantan have 2.232 times made ≥4 ANC visits compared to women in the Papua region. Women in Sulawesi have 1,980 times more than AN4 ANC visits compared to women in the Papua region. In addition to the region category, other variables found to contribute to the predictor were age, husband/partner, education, parity, wealth and insurance.

**Conclusion:** There were disparities between regions in the ANC utilization in Indonesia.

## Introduction

Indonesia has entered the final year of the 2015-2019 National Medium-Term Development Plan. In the 2015-2019 National Medium-Term Development Plan, 4 main health targets that must be achieved by 2019 have been established, namely: 1) Improving the health and nutritional status of the community; 2) Improve control of communicable and non-communicable diseases; 3) Increase the equity and quality of health services; 4) Increase financial protection, availability, distribution, quality of medicines and health resources [1].

In health development, the target of increasing equal distribution and quality of health services is determined by three indicators, namely the number of sub-districts that have at least one accredited Puskesmas (Health Center) of 5,600, the number of regencies/cities that have at least one nationally accredited hospital of 481, and the percentage of regencies/cities that have up to 80% complete basic immunization in infants as much as 95%. Based on the Ministry of Health’s report the target has been achieved, the target number of sub-districts that have at least one Health Center accredited in 2018 of 4900 sub-districts, has been realized as many as 5,385 Sub-districts (109.9%), or around 7,518 Health Centers. This achievement exceeded the target set because several regencies/cities used the Regional Revenue and Expenditure Budget purely for the accreditation process, not from the Non-Physical Allocation Fund. For the number of regencies/cities that have at least one nationally accredited hospital, the realization in 2018 was 440 (101.4%) of the target of 434. For the immunization target still not achieved, 2018 data shows complete basic immunization coverage for children aged 12-23 months in Indonesia is 57.9%, incomplete is 32.9% and not immunized is 9.2% [2].

In the target of improving the community’s health and nutrition status, several achievement targets have been set, namely the maternal mortality rate (MMR) of 306/100,000 live births, the infant mortality rate (IMR) which is targeted to reach 24/1000 live births, the prevalence of malnutrition in children under five 17/100,000, and the prevalence of stunting in children under two years 28/100,000 population. MMR is currently reported to have decreased by 346 deaths to 305 maternal deaths per 100,000 live births but has not reached the MDG target in 2015 of 102/100,000 live births [3]. On the other hand, Indonesia is demanded higher on the SDG’s target to reduce MMR to below 70/100,000 live births and reduce neonatal mortality to 12/1000 live births and Toddler Death Rate 25/1000 live births [4]. MMR in Indonesia is the highest compared to other ASEAN countries which are 9 times compared to Malaysia, 5 times that of Vietnam and almost 2 times that of Cambodia. Based on WHO reports, the estimated MMR in developed countries is 12/100,000 live births, while in developing countries it is 239/100,000 live births [5–6].

The results of research in Indonesia that used 2013 data showed disparities in maternal deaths among districts/cities in Indonesia, with the highest risk of maternal deaths occurring in Eastern Indonesia. The risk factors that most influenced maternal mortality were population density with OR 0.283 (95% CI 0.185-0.430) and delivery by health workers with OR 1.745 (95% CI 1.081-2.815). The risk of maternal death is high in districts/cities with low coverage of fourth pregnancy visit, low coverage of delivery by health workers, low postpartum visit coverage, the high average number of children, average length of schooling for women of childbearing age, and poverty high [7].

The main strategy for decreasing maternal morbidity and mortality with antenatal care (ANC). ANC aims to monitor and maintain the health and safety of the mother and fetus, detect all complications of pregnancy and take the necessary actions, respond to complaints, prepare for birth, and promote healthy living behavior. ANC visits are very important to detect and prevent unwanted things that arise during pregnancy [8]. In developing countries, there has been an increase in the utilization of maternal health services but it still varies among population groups. Disparities can occur due to geographical, demographic, socioeconomic, and cultural differences. Gaps that occur result in decreased access to services, service quality, and service affordability [9–10].

In 2018 there has been an increase in the proportion of ANC visits to women aged 10-54 years ie first visit by 96.1% compared to 2013 by 95.2%, while for ANC fourth visits in 2018 amounted to 74.1% compared to 2013 by 70.0%, the coverage of ANC fourth visits is still below the target set in the 2017 Strategic Plan which is 76.0% [11]. However, the quality of services to ensure early diagnosis and appropriate care for pregnant women still needs to be improved. Midwives spearheading to serve pregnancy checks including identification of complications or symptoms of complications, assist in labor and childbirth examination. If there are signs of complications that cannot be treated, the midwife must make a referral to a health facility that provides Basic Emergency Neonatal Obstetric Services, to obtain further treatment [12]. Data from the Ministry of Health in 2018 stated that the majority (62.7%) of deliveries were assisted by midwives and were carried out in independent midwife practices (29%), although there were still many at home (16%) [11].

This study was conducted to analyze interregional disparities in the utilization of ≥4 ANC visits during pregnancy in women aged 15-49 years who gave birth in the last five years in Indonesia. This study is important to do so that it can provide clear directions for the Ministry of Health to complete regional priority data in an effort to reduce maternal mortality.

## Methods

### Data Source

This study is an analysis of the 2017 Indonesian Demographic Data Survey (IDHS) data. The IDHS was part of the International Demographic and Health Survey (DHS) program conducted by the Inner City Fund (ICF) which uses stratification and multistage random sampling methods. The unit of analysis in this study is women aged 15-49 years old who had given birth in the last 5 years, to obtain a sample of 15,351 women.

### Procedure

Ethical clearance has been obtained in the 2017 IDHS from the National Ethics Committee. The respondents’ identities have all been deleted from the dataset. Respondents have provided written approval for their involvement in the study. Through the website: https://dhsprogram.com/data/new-user-registration.cfm researchers have obtained permission to use data for the purposes of this study.

### Data Analysis

The Ministry of Health of the Republic of Indonesia recommends that the ANC during pregnancy be done at least 4 times, namely in the first trimester 1 time, in the second trimester 1 time, and in the third trimester 2 times [13]. Other variables analyzed as independent variables are the place of residence, age, husband/partner, education level, parity, wealth status, and health insurance. Because all variables are categorical, the Chi-square test was used to select variables related to the frequency of ANC utilization during pregnancy. Because of the nature of the dependent variable, Binary Logistic Regression is used for the final test to determine disparity. SPSS 21 software is used for all stages of statistical analysis.

## Results

Fig 1 is a description of the distribution of ≥4 ANC visits in 34 provinces in Indonesia. Seen in the East (Maluku and Papua Region) has the lowest distribution of ≥4 ANC visits. While in the westernmost region (part of Sumatra Region) it has a distribution of ≥4 ANC visits one level above. While the distribution of ≥4 ANC visits is best centered in the central region of the Java-Bali Region.

**Fig 1.**
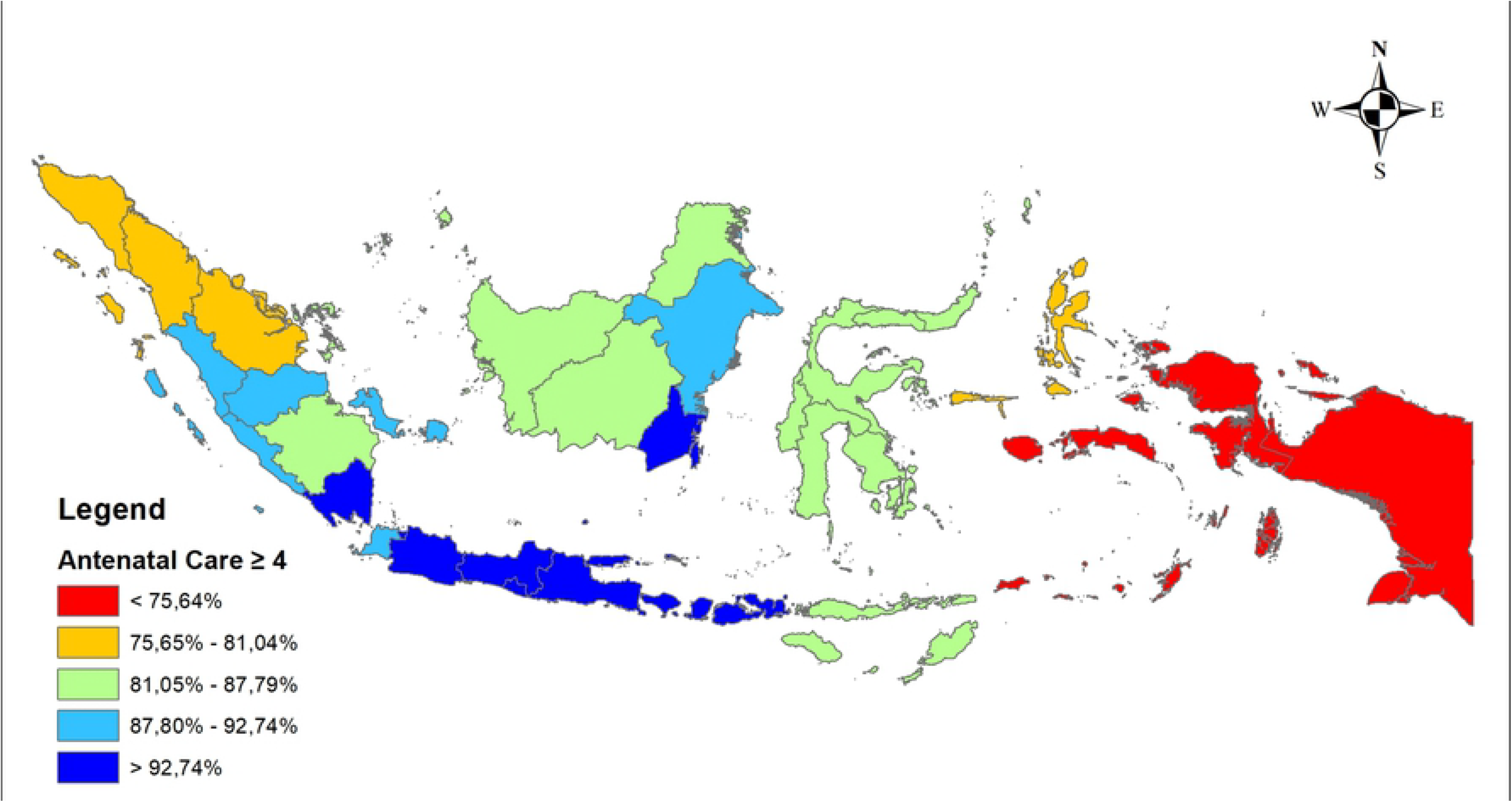
Distribution of ≥4 ANC visits by provinces in Indonesia.

The statistical description of female respondents aged 15-49 years who gave birth in the last five years in Indonesia is presented in Table 1. Table 1 shows that there are statistically significant differences between regions. Each region was seen dominated by the use of ANC which had ≥4 visits.

**Table 1.** Socio-Demographic of Respondents (n=15,351)

**Tabel 1. Socio-Demographic of Respondents (n=15,351)**
| Variables | Region |  |  |  |  |  |  | All | P |
| --- | --- | --- | --- | --- | --- | --- | --- | --- | --- |
|  | Sumatera | Java-Bali | Nusa Tenggara | Kalimantan | Sulawesi | Maluku Islands | Papua |  |  |
| ANC |  |  |  |  |  |  |  |  | 0.000*** |
| • <4 (ref.) | 606 (15.07%) | 261 (5.36%) | 120 (9.30%) | 154 (10.82%) | 310 (13.53%) | 248 (24.17%) | 121 (27.94%) | 1820 (11.86%) |  |
| • ≥4 | 3416 (84.93%) | 4605 (94.64%) | 1170 (90.70%) | 1269 (89.18%) | 1981 (86.47%) | 778 (75.83%) | 312 (72.06%) | 13531 (88.14%) |  |
| Place of Residence |  |  |  |  |  |  |  |  | 0.000*** |
| • Urban | 1807 (44.93%) | 3312 (68.06%) | 384 (29.77%) | 738 (51.86%) | 841 (36.71%) | 380 (37.04%) | 106 (24.48%) | 7568 (49.30%) |  |
| • Rural (ref.) | 2215 (55.07%) | 1554 (31.94%) | 906 (70.23%) | 685 (48.14%) | 1450 (63.29%) | 646 (62.96%) | 327 (75.52%) | 7783 (50.70%) |  |
| Age group of respondents |  |  |  |  |  |  |  |  | 0.000*** |
| • 15-19 | 96 (2.39%) | 102 (2.10%) | 32 (2.48%) | 46 (3.23%) | 75 (3.27%) | 51 (4.97%) | 14 (3.23%) | 416 (2.71%) |  |
| • 20-24 | 544 (13.53%) | 791 (16.26%) | 199 (15.43%) | 231 (16.23%) | 420 (18.33%) | 154 (15.01%) | 75 (17.32%) | 2414 (15.73%) |  |
| • 25-29 | 1005 (24.99%) | 1222 (25.11%) | 317 (24.57%) | 390 (27.41%) | 555 (24.23%) | 239 (23.29%) | 119 (27.48%) | 3847 (25.06%) |  |
| • 30-34 | 1146 (28.49%) | 1226 (25.20%) | 328 (25.43%) | 366 (25.72%) | 534 (23.31%) | 260 (25.34%) | 103 (23.79%) | 3963 (25.82%) |  |
| • 35-39 | 829 (20.61%) | 1021 (20.98%) | 254 (19.69%) | 241 (16.94%) | 429 (18.73%) | 199 (19.40%) | 83 (19.17%) | 3056 (19.91%) |  |
| • 40-44 | 340 (8.45%) | 419 (8.61%) | 127 (9.84%) | 115 (8.08%) | 233 (10.17%) | 93 (9.06%) | 30 (6.93%) | 1357 (8.84%) |  |
| • 45-49 (ref.) | 62 (1.54%) | 85 (1.75%) | 33 (2.56%) | 34 (2.39%) | 45 (1.96%) | 30 (2.93%) | 9 (2.08%) | 298 (1.94%) |  |
| Have a husband/partner |  |  |  |  |  |  |  |  | 0.000*** |
| • No (ref.) | 126 (3.13%) | 135 (2.77%) | 74 (5.74%) | 44 (3.09%) | 71 (3.10%) | 33 (3.22%) | 25 (5.77%) | 508 (3.31%) |  |
| • Yes | 3896 (96.87%) | 4731 (97.23%) | 1216 (94.26%) | 1379 (96.91%) | 2220 (96.90%) | 993 (96.78%) | 408 (94.23%) | 14843 (96.69%) |  |
| Education Level |  |  |  |  |  |  |  |  | 0.000*** |
| • No education (ref.) | 38 (0.94%) | 21 (0.43%) | 56 (4.34%) | 14 (0.98%) | 36 (1.57%) | 8 (0.78%) | 31 (7.16%) | 204 (1.33%) |  |
| • Primary | 888 (2.08%) | 1185 (24.35%) | 439 (34.03%) | 404 (28.39%) | 630 (27.50%) | 224 (21.83%) | 89 (20.55%) | 3859 (24.14%) |  |
| • Secondary | 2297 (57.11%) | 2979 (61.22%) | 594 (46.05%) | 795 (55.87%) | 1156 (50.46%) | 578 (56.34%) | 229 (52.89%) | 8628 (56.20%) |  |
| • Higher | 799 (19.87%) | 681 (14.00%) | 201 (15.58%) | 210 (14.76%) | 469 (20.47%) | 216 (21.05%) | 84 (19.40%) | 2660 (17.33%) |  |
| Parity |  |  |  |  |  |  |  |  | 0.000*** |
| • < 2 | 1161 (28.87%) | 1758 (36.13%) | 370 (28.68%) | 401 (28.18%) | 694 (30.29%) | 267 (26.02%) | 104 (24.02%) | 4755 (30.98%) |  |
| • 2 - 4 | 2572 (63.95%) | 2949 (60.60%) | 757 (58.68%) | 936 (65.78%) | 1362 (59.45%) | 588 (57.31%) | 243 (56.12%) | 9407 (61.28%) |  |
| • > 4 (ref.) | 289 (7.19%) | 159 (3.27%) | 163 (12.64%) | 86 (6.04%) | 235 (10.26%) | 171 (16.67%) | 86 (19.86%) | 1189 (7.75%) |  |
| Wealth status |  |  |  |  |  |  |  |  | 0.000*** |
| • Poorest (ref.) | 871 (21.66%) | 474 (9.74%) | 783 (60.70%) | 285 (20.03%) | 858 (37.45%) | 576 (56.14%) | 226 (52.19%) | 4073 (25.53%) |  |
| • Poorer | 876 (21.78%) | 805 (16.54%) | 244 (18.91%) | 317 (22.28%) | 517 (22.57%) | 193 (18.81%) | 79 (18.24%) | 3031 (19.74%) |  |
| • Midle | 857 (21.31%) | 1050 (21.58%) | 118 (9.15%) | 342 (24.03%) | 351 (15.32%) | 117(11.40%) | 55 (12.70%) | 2890 (18.83%) |  |
| • Richer | 752 (18.70%) | 1259 (25.87%) | 77 (5.97%) | 254 (17.85%) | 272 (11.87%) | 103 (10.04%) | 43 (9.93%) | 2760 (17.98%) |  |
| • Richest | 666 (16.56%) | 1278 (26.26%) | 68 (5.27%) | 225 (15.81%) | 293 (12.79%) | 37 (3.61%) | 30 (6.93%) | 2597 (16.92%) |  |
| Covered by health insurance |  |  |  |  |  |  |  |  | 0.000*** |
| • No (ref.) | 1455 (36.18%) | 1961 (40.30%) | 499 (38.68%) | 630 (44.27%) | 712 (31.08%) | 482 (46.98%) | 100 (23.09%) | 5839 (38.04%) |  |
| • Yes | 2567 (63.82%) | 2905 (59.70%) | 791 (61.32%) | 793 (55.73%) | 1579 (68.92%) | 544 (53.02%) | 333 (76.91%) | 9512 (61.96%) |  |

Note: \* p < 0.05; \*\* p < 0.01; \*\*\* p < 0.001.

Table 1 informs that in the Java-Bali and Kalimantan regions it is more dominated by urban areas, while the remaining regions are dominated by rural areas. In all regions, it was also seen that women were dominated in the 25-29 year and 30-34 year age categories. Table 1 shows that in all regions dominated by women who have a husband/partner, have a secondary education level, and have 2-4 parity.

Table 1 shows that almost all regions are dominated by women who have the wealth status of the poorer or poorest, except in the Java-Bali region which is more dominant in women with the richest wealth status. Women aged 15-49 who had delivered their last five years in Indonesia were dominantly covered by health insurance in all regions.

Table 2 shows the results of the binary logistic regression test which shows disparities between regions in the use of ANC in Indonesia. At this stage <4 ANC visits during pregnancy are used as a reference. Table 3 shows that all regions show gaps with the Papua region as a reference, except the Maluku region which is not significant shows differences in the use of ANC with the Papua region.

**Table 2.** Binary Logistic Regression of ANC Utilization (n=15,351)

| Predictor | ≥4 ANC visits |  |  |  |
| --- | --- | --- | --- | --- |
|  | P | OR | Lower Bound | Upper Bound |
| Region: Sumatera | 0.014* | 1.370 | 1.066 | 1.761 |
| Region: Java-Bali | 0.000*** | 3.607 | 2.741 | 4.746 |
| Region: Nusa Tenggara | 0.000*** | 4.365 | 3.229 | 5.899 |
| Region: Kalimantan | 0.000*** | 2.232 | 1.664 | 2.994 |
| Region: Sulawesi | 0.000*** | 1.980 | 1.523 | 2.574 |
| Region: Maluku Islands | 0.171 | 1.213 | 0.920 | 1.600 |
| Place of Residence: Urban | 0.584 | 0.967 | 0.856 | 1.092 |
| Age group of respondents: 15-19 | 0.000*** | 0.336 | 0.218 | 0.519 |
| Age group of respondents: 20-24 | 0.038* | 0.675 | 0.465 | 0.979 |
| Age group of respondents: 25-29 | 0.691 | 0.930 | 0.652 | 1.328 |
| Age group of respondents: 30-34 | 0.763 | 1.055 | 0.744 | 1.496 |
| Age group of respondents: 35-39 | 0.441 | 1.145 | 0.811 | 1.618 |
| Age group of respondents: 40-44 | 0.841 | 1.037 | 0.725 | 1.485 |
| Have a husband/partner: Yes | 0.000*** | 2.107 | 1.674 | 2.651 |
| Education Level: Primary | 0.000*** | 2.527 | 1.838 | 3.474 |
| Education Level: Secondary | 0.000*** | 3.882 | 2.815 | 5.353 |
| Education Level: Higher | 0.000*** | 3.669 | 2.559 | 5.259 |
| Parity: < 2 | 0.000*** | 3.580 | 2.857 | 4.485 |
| Parity: 2 - 4 | 0.000*** | 2.519 | 2.121 | 2.992 |
| Wealth status: Poorer | 0.000*** | 1.674 | 1.449 | 1.933 |
| Wealth status: Middle | 0.000*** | 2.056 | 1.739 | 2.431 |
| Wealth status: Richer | 0.000*** | 2.690 | 2.204 | 3.284 |
| Wealth status: Richest | 0.000*** | 3.596 | 2.813 | 4.596 |
| Covered by health insurance: Yes | 0.000*** | 1.485 | 1.334 | 1.653 |

Note: \* p < 0.05; \*\* p < 0.01; \*\*\* p < 0.001.

Table 2 shows that the biggest gap in the utilization of ≥4 ANC visits is between the Nusa Tenggara and Papua regions. Women in the Nusa Tenggara region have 4.365 times more than ≥4 ANC visits compared to women in the Papua region (OR 4.365; 95% CI 3.229-5.899). While women in the Java-Bali region were 3.607 times more likely to make ≥4 ANC visits than women in the Papua region (OR 3.607; 95% CI 2.741-4.746).

Table 2 also shows disparities between the Sumatra, Kalimantan and Sulawesi regions compared to the Papua region. Women in the Sumatra region have a 1.370 chance of making ≥4 ANC visits compared to women in the Papua region (OR 1.370; 95% CI 1.066-1.761). Women in the Kalimantan region had 2.232 times made ≥4 ANC visits compared to women in the Papua region (OR 2.232; 95% CI 1.664-2.994). Women in the Sulawesi region had 1,980 chances of making ≥4 ANC visits compared to women in the Papua region (OR 1.980; 95% CI 1.523-2.574).

In addition to the region category, other variables found to contribute to the predictor are age group, husband/partner, education level, parity, wealth status, and health insurance. Table 2 shows that women in the 15-19 age group had 0.336 times more than ≥4 ANC visits compared to women in the 45-49 age group (OR 0.336; 95% CI 0.218-0.519). While the 20-24 years age group had a 0.675 chance of doing ≥4 ANC visits compared to women in the 45-49 age group (OR 0.675; 95% CI 0.465-0.979). This shows that the very young age group has a lower possibility than the oldest age group.

Table 2 informs that women who have a husband/partner have a better chance of making ≥4 ANC visits than those without. Women who have a husband/partner have 2.107 times more than AN4 ANC visits compared to women who do not have a husband/partner (OR 2.107; 95% CI 1.674-2.651).

Table 2 shows that women with better levels of education have a better chance of making ≥4 ANC visits than those without education. Women with primary education have 2.527 chances of doing ≥4 ANC visits compared to women with no education (OR 2.527; 95% CI 1.838-3.474). Women with secondary education were 3.882 times more than ≥4 ANC visits compared to women with no education (OR 3.882; 95% CI 2.815-5.353). Women with a higher level of education had a 3.669 times chance ≥4 ANC visits than women with no education (OR 3.669; 95% CI 2.559-5.259).

Table 2 shows that women with lower parity have a better chance of making ≥4 ANC visits than those who have parity >4. Women who have parity <2 have a 3.580 times chance of doing AN4 ANC visits than women who have parity >4 (OR 3.580; 95% CI 2.857-4.485). Women who had parity 2-4 had 2.519 times more than ≥4 ANC visits compared to women who had parity >4 (OR 2.519; 95% CI 2.121-2.992).

Table 2 informs us that the better the wealth status held by a woman, the higher the probability of making ≥4 ANC visits. richest women had 3.596 chances of making ≥4 ANC visits compared to the poorest women (OR 3.596; 95% CI 2.813-4.586).

Table 2 shows that women covered by health insurance had a better chance of making to do ≥4 ANC visits than those who were not covered. Women who are covered by health insurance are likely to get 1.485 times ≥4 ANC visits compared to women who are not covered by health insurance (OR 1.485; 95% CI 1.334-1.653).

## Discussion

The results showed that disparity between regions in the use of ANC is still ongoing. The disparity is also clearly seen between the East and West regions. The results of this analysis are in line with several studies in Indonesia which show that the Eastern region is lagging behind the Western region [14–16]. Especially when compared to the Java-Bali region as the center of government.

Geographically, conditions in Eastern Indonesia also show more extreme variability than Western regions. This condition makes some parts of the East in the category of isolated or remote area [17–18], and some other areas are quite difficult to reach because of the limited means of roads and public transportation available [19].

Qualitatively, some research also shows that in the Eastern region having more health beliefs is a challenge for health workers to strive for better maternal health [20–21]. Not only applies to the community, but the health belief also encompasses health workers, because they are an inseparable part of the community itself [22].

The analysis shows that there is no difference between urban and rural areas in ANC utilization in Indonesia. This condition is different from the findings in Nigeria [23], Ethiopia [24], Pakistan [25] and several other countries [26], which found disparities between urban and rural areas.

The age group was found to be a predictor of ANC utilization. The youngest age group has a lower probability of making ≥4 ANC visits. This is likely due to the lack of experience, so knowledge about health risks is lower [27–28]. A study in India that analyzed the relationship between child marriage and the utilization of maternal healthcare services concluded that many challenges were found so that more effort was needed so that child marriage could have a positive impact on the use of maternal healthcare services [29].

The analysis shows that women who have husbands/partners are likely to be better at using ANC. This is in line with the conclusions in several studies that show the role of husband/partner in providing support for a woman’s health behavior [30–33]. Some other studies actually involve a husband to be able to improve a woman’s health status through actively better health behaviors [34–35].

Analysis of this study proves that education is one of the determining factors for women in Indonesia to make ≥4 ANC visits. In general, it can be explained that the more educated a person is, the easier it is to receive new health information, while at the same time being able to understand the dangers or risks of behaviors that have an impact on health [36–38]. Education has also been shown to play a role in one’s perception of the quality of health services [39, 40]. Furthermore, improving education is generally accepted as one of the determinants of life expectancy [41].

This study found that parity is a determinant of the use of ANC. The lower the parity, the more likely it is to make ≥4 ANC visits. Parity as one of the determinants of ANC utilization is also found in several recent studies in several countries [42–44].

In line with the level of education, wealth status was also found to be directly proportional to the behavior of ≥4 ANC visits. This result is in accordance with several studies which found that wealth status is one of the positive determinants of ANC utilization, namely in Ethiopia [45], Pakistan [46], Nigeria [47], and Uganda [48]. The better the wealth status of a woman, the more likely it is to make ≥4 ANC visits.

Women covered by health insurance were found to have better use of ANC. Women who do not have health insurance have lower ANC utilization. This finding is in line with the goal of the National Health Insurance released by the Indonesian government to provide universal access to health care facilities [49, 50]. Social insurance policies to increase public access to health care facilities have also been adopted by other countries. The results of studies evaluating this matter show positive results [51–53], although also in the implementation there were still some obstacles encountered [54–55].

Disparities found and detected in this study are still limited to superficial. Researchers suggest further research in order to detect the causes of disparity in more depth.

## Conclusions

Based on the results of the study it can be concluded that there are 10 variables that become a barrier for Indonesian women to make AN4 ANC visits during pregnancy. The barriers consisted of variables of young age, low education, high parity, poverty, not having health insurance, not being able to read, not being exposed to the media, never using the internet, not knowing the danger signs of pregnancy, and belief in traditional birth attendants. Thus, maternal health programs need to address barriers to things for effective health care utilization.

## Acknowledgments

The author would like to thank the ICF International, who has agreed to allow the 2017 IDHS data to be analyzed in this article.

## Author Contributions

Conceptualization: Agung Dwi Laksono.

Data curation: Agung Dwi Laksono, Ratna Dwi Wulandari.

Formal analysis: Ratna Dwi Wulandari.

Methodology: Agung Dwi Laksono, Rukmini Rukmini.

Writing ± original draft: Agung Dwi Laksono, Rukmini Rukmini.

Writing ± review & editing: Ratna Dwi Wulandari.

## References

1. Ministry of Health. Decree of the Minister of Health of the Republic of Indonesia Number HK 02.02 / MENKES / 52/2015 about Ministry of Health Strategic Plan for 2015-2019. HK 02.02 / MENKES / 52/2015 Indonesia; 2015.

2. Directorate General of Health Services. The 2018 Directorate General of Health Services Performance Accountability Report [Internet]. Jakarta; 2019. Available from: http://yankes.kemkes.go.id/app/lakip2/downloads/2017/KP/ditjen/lakip_ditjen_2017.pdf

3. Data and Information Center Ministry of Health. Mother’s Day: Maternal Health Situation [Internet]. Jakarta; 2014. Available from: http://www.depkes.go.id/download.php?file=download/pusdatin/infodatin/infodatin-ibu.pdf

4. Communication and Community Service Bureau Ministry of Health. 4 Health Targets Must Be Achieved by 2019 (4 Target Kesehatan ini Harus Tercapai di 2019) [Internet]. Press Release. 2019. p. 1–4. Available from: http://www.depkes.go.id/article/view/18030700008/4-target-kesehatan-ini-harus-tercapai-di-2019.html

5. World Health Organization. Trends in maternal mortality: 1990 to 2015: estimates by WHO, UNICEF, UNFPA, World Bank Group and the United Nations Population Division. [Internet]. Geneva; 2015. Available from: https://apps.who.int/iris/bitstream/handle/10665/194254/9789241565141_eng.pdf;jsessionid=AB201B62E0913D576E5CF9430F900F98?sequence=1

6. Achadi EL. Maternal and Neonatal Death in Indonesia (Kematian Maternal dan Neotatal di Indonesia) [Internet]. Jakarta; 2019. Available from: http://www.depkes.go.id/resources/download/info-terkini/rakerkesnas-2019/SESII/Kelompok1/1-Kematian-Maternal-dan-Neonatal-di-Indonesia.pdf

7. Nurrizka RH, Wahyono TYM. Disparity of Maternal Mortality in Indonesia: Ecological Study with Spatial Analysis. Media Kesehat Masy Indones. 2018;14(2):119–27. https://doi.org/10.30597/mkmi.v14i2.3630.

8. Hijazi HH, Alyahya MS, Sindiani AM, Saqan RS, Okour AM. Determinants of antenatal care attendance among women residing in highly disadvantaged communities in northern Jordan: a cross-sectional study. Reprod Health. 2018;15(1):Article number 106. https://doi.org/10.1186/s12978-018-0542-3.

9. Bobo FT, Yesuf EA, Woldie M. Inequities in utilization of reproductive and maternal health services in Ethiopia. Int J Equity Health. 2017;16(1):Article number 105. https://doi.org/10.1186/s12939-017-0602-2.

10. Chi PC, Bulage P, Urdal H, Sundby J. A qualitative study exploring the determinants of maternal health service uptake in post-conflict Burundi and Northern Uganda. BMC Pregnancy Childbirth. 2015;15(1):Article number 18. https://doi.org/10.1186/s12884-015-0449-8.

11. National Institute of Health Research and Development of Ministry of Health of the Republic of Indonesia. The 2018 Indonesia Basic Health Survey (Riskesdas): National Report. Jakarta; 2019.

12. Ministry of National Development Planning / National Development Planning Agency. Collection of Sectoral Study and Evaluation Summary 2008-2013. Accelerate Maternal Mortality Rate [Internet]. Jakarta; 2014. Available from: https://www.bappenas.go.id/files/ekps/2014/4.KumpulanRingkasanKajiandanEvaluasiSektoral2008-2013.pdf

13. National Population and Family Planning Board, Statistics Indonesia, Ministry of Health, The DHS Program. Indonesia Demographic and Health Survey 2017 [Internet]. Jakarta; 2018. Available from: https://www.dhsprogram.com/pubs/pdf/FR342/FR342.pdf

14. Mubasyiroh R, Nurhotimah E, Laksono AD. Health Service Accessibility Index in Indonesia (Indeks Aksesibilitas Pelayanan Kesehatan di Indonesia). In: Supriyanto S, Chalidyanto D, Wulandari RD, editors. Accessibility of Health Services in Indonesia (Aksesibilitas Pelayanan Kesehatan di Indonesia). Jogjakarta: PT Kanisius; 2016. p. 21–58.

15. Yudhistira MH, Sofiyandi Y. Seaport status, port access, and regional economic development in Indonesia. Marit Econ Logist. 2018;20(4):549–68. https://doi.org/10.1057/s41278-017-0089-1.

16. Laksono AD, Wulandari RD, Soedirham O. Regional Disparities of Health Center Utilization in Rural Indonesia. Malaysian J Public Heal Med. 2019;19(1).

17. Suharmiati, Laksono AD, Astuti WD. Policy Review on Health Services in Primary Health Center in the Border and Remote Area (Review Kebijakan tentang Pelayanan Kesehatan Puskesmas di Daerah Terpencil Perbatasan). Bull Heal Syst Res. 2013;16(2):109–16.

18. United Nations Group of Experts on Geographical Names. United Nations Conference on the Standardization of Geographical Names, 11th [Internet]. 2017 [cited 2018 Sep 1]. Available from: https://unstats.un.org/unsd/geoinfo/UNGEGN/ungegnConf11.html

19. Soewondo P, Johar M, Pujisubekti R, Halimah H, Irawati DO. INSPECTING PRIMARY HEALTHCARE CENTERS IN REMOTE AREAS: FACILITIES, ACTIVITIES, AND FINANCES. J Adm Kesehat Indones. 2019;7(1):89–98. https://doi.org/10.20473/jaki.v7i1.2019.89-98.

20. Dwiningsih S, Laksono AD. How to control the sexually transmitted diseases in Benjina?: qualitative studies on the practice of prostitution. Heal Sci J Indones. 2019;10(1):58–66. https://doi.org/10.22435/hsji.v10i1.1044.

21. Laksono AD, Soerachman R, Angkasawati TJ. Case Study of Muyu Ethnic’s Maternal Health in Mindiptara District-Boven Digoel (Studi Kasus Kesehatan Maternal Suku Muyu di Distrik Mindiptana, Kabupaten Boven Digoel). J Reprod Heal. 2016;07/03:145–55. https://doi.org/10.22435/kespro.v7i3.4349.145-155.

22. Laksono AD, Faizin K. Traditions Influence Into Behavior in Health Care; Ethnographic Case Study on Health Workers Muyu Tribe. Bull Heal Syst Res. 2015;18(4):347–54. https://doi.org/10.22435/hsr.v18i4.4567.347-354.

23. Adewuyi EO, Auta A, Khanal V, Bamidele OD, Akuoko CP, Adefemi K, et al. Prevalence and factors associated with underutilization of antenatal care services in Nigeria: A comparative study of rural and urban residences based on the 2013 Nigeria demographic and health survey. PLoS One. 2018;13(5):Article number e0197324. https://doi.org/10.1371/journal.pone.0197324

24. Bobo FT, Yesuf EA, Woldie M. Inequities in utilization of reproductive and maternal health services in Ethiopia. Int J Equity Health. 2017;16(9). doi: 10.1186/s12939-017-0602-2. https://doi.org/10.1371/journal.pone.0197324

25. Sahito A, Fatmi Z. Inequities in antenatal care, and individual and environmental determinants of utilization at national and sub-national level in Pakistan: A multilevel analysis. Int J Heal Policy Manag. 2018;7(8):699–710. https://doi.org/10.1371/journal.pone.0197324

26. Sully EA, Biddlecom AS, Darroch JE. Not all inequalities are equal: Differences in coverage across the continuum of reproductive health services. BMJ Glob Heal. 2019;4(5):Article number e001695. https://doi.org/10.1371/journal.pone.0197324

27. Hattar-Pollara M. Barriers to Education of Syrian Refugee Girls in Jordan: Gender-Based Threats and Challenges. J Nurs Scholarsh. 2019;51(3):241–51. https://doi.org/10.1111/jnu.12480.

28. Dey A, Hay K, Afroz B, Chandurkar D, Singh K, Dehingia N, et al. Understanding intersections of social determinants of maternal healthcare utilization in Uttar Pradesh, India. PLoS One. 2018;13(10):Article number e0204810. https://doi.org/10.1371/journal.pone.0204810.

29. Paul P, Chouhan P. Association between child marriage and utilization of maternal health care services in India: Evidence from a nationally representative cross-sectional survey. Midwifery. 2019;75:66–71. https://doi.org/10.1016/j.midw.2019.04.007.

30. Blanchard AK, Nair SG, Bruce SG, Ramanaik S, Thalinja R, Murthy S, et al. A community-based qualitative study on the experience and understandings of intimate partner violence and HIV vulnerability from the perspectives of female sex workers and male intimate partners in North Karnataka state, India. BMC Womens Health. 2018;18(1):Article number 66. https://doi.org/10.1186/s12905-018-0554-8.

31. Sumankuuro J, Mahama MY, Crockett J, Wang S, Young J. Narratives on why pregnant women delay seeking maternal health care during delivery and obstetric complications in rural Ghana. BMC Pregnancy Childbirth. 2019;19(1):Article number 260. https://doi.org/10.1186/s12884-019-2414-4.

32. Jungari S, Paswan B. What he knows about her and how it affects her? Husband’s knowledge of pregnancy complications and maternal health care utilization among tribal population in Maharashtra, India. BMC Pregnancy Childbirth. 2019;19(1):Article number 70. https://doi.org/10.1186/s12884-019-2214-x.

33. Sakuma S, Yasuoka J, Phongluxa K, Jimba M. Determinants of continuum of care for maternal, newborn, and child health services in rural Khammouane, Lao PDR. PLoS One. 2019;14(4):Article number e0215635. https://doi.org/10.1371/journal.pone.0215635.

34. Baraki Z, Wendem F, Gerensea H, Teklay H. Husbands involvement in birth preparedness and complication readiness in Axum town, Tigray region, Ethiopia, 2017. BMC Pregnancy Childbirth. 2019;19(1):Article number 180. https://doi.org/10.1186/s12884-019-2338-z.

35. Ahmed S, Jafri H, Rashid Y, Yi H, Dong D, Zhu J, et al. Autonomous decision-making for antenatal screening in Pakistan: views held by women, men and health professionals in a low–middle income country. Eur J Hum Genet. 2019;27(6). https://doi.org/10.1038/s41431-019-0353-1.

36. Jafaralilou H, Zareban I, Hajaghazadeh M, Matin H, Didarloo A. The impact of theory-based educational intervention on improving helmet use behavior among workers of cement factory, Iran. J Egypt Public Health Assoc. 2019;94(1):Article number 1. https://doi.org/10.1186/s42506-018-0001-6.

37. Ba DM, Ssentongo P, Agbese E, Kjerulff KH. Prevalence and predictors of contraceptive use among women of reproductive age in 17 sub-Saharan African countries: A large population-based study. Sex Reprod Healthc. 2019;21:26–32. https://doi.org/10.1186/s42506-018-0001-6.

38. Teye-Kwadjo E. Risky driving behaviour in urban Ghana: the contributions of fatalistic beliefs, risk perception, and risk-taking attitude. Int J Heal Promot Educ. 2019;57(5):256–73. https://doi.org/10.1080/14635240.2019.1613163.

39. Påfs J, Musafili A, Binder-Finnema P, Klingberg-Allvin M, Rulisa S, Essén B. Beyond the numbers of maternal near-miss in Rwanda - a qualitative study on women’s perspectives on access and experiences of care in early and late stage of pregnancy. BMC Pregnancy Childbirth. 2016;16(1):Article number 257. https://doi.org/10.1186/s12884-016-1051-4.

40. Megatsari H, Laksono AD, Ridlo IA, Yoto M, Azizah AN. Community Perspective about Health Services Access. Bul Penelit Sist Kesehat. 2018;21:247–253. https://doi.org/10.1186/s12884-016-1051-4.

41. Luy M, Zannella M, Wegner-Siegmundt C, Minagawa Y, Lutz W, Caselli G. The impact of increasing education levels on rising life expectancy: a decomposition analysis for Italy, Denmark, and the USA. Genus. 2019;75(1):Article number 11. https://doi.org/10.1186/s41118-019-0055-0.

42. You H, Yu T, Gu H, Kou Y, Xu X-P, Li X-L, et al. Factors Associated With Prescribed Antenatal Care Utilization: A Cross-Sectional Study in Eastern Rural China. Inq (United States). 2019;56. https://doi.org/10.1177/0046958019865435.

43. Tikmani SS, Ali SA, Saleem S, Bann CM, Mwenechanya M, Carlo WA, et al. Trends of antenatal care during pregnancy in low- and middle-income countries: Findings from the global network maternal and newborn health registry. Semin Perinatol. 2019;43(5):297–307. https://doi.org/10.1053/j.semperi.2019.03.020.

44. Mumtaz S, Bahk J, Khang Y-H. Current status and determinants of maternal healthcare utilization in Afghanistan: Analysis from Afghanistan demographic and health survey 2015. PLoS One. 2019;14(6):Article number e0217827. https://doi.org/10.1371/journal.pone.0217827.

45. Mekonnen T, Dune T, Perz J, Ogbo FA. Trends and determinants of antenatal care service use in ethiopia between 2000 and 2016. Int J Environ Res Public Health. 2019;16(5):Article number 748. https://doi.org/10.3390/ijerph16050748.

46. Zakar R, Zakar MZ, Aqil N, Chaudhry A, Nasrullah M. Determinants of maternal health care services utilization in Pakistan: evidence from Pakistan demographic and health survey, 2012-13. J Obstet Gynaecol (Lahore). 2017;37(3):330–7. https://doi.org/10.1080/01443615.2016.1250728.

47. Olaitan T, Okafor IP, Onajole AT, Abosede OA. Ending preventable maternal and child deaths in western Nigeria: Do women utilize the life lines? PLoS One. 2017;12(5):Article number e0176195. https://doi.org/10.1371/journal.pone.0176195.

48. Wilson M, Patterson K, Nkalubo J, Lwasa S, Namanya D, Twesigomwe S, et al. Assessing the determinants of antenatal care adherence for Indigenous and non-Indigenous women in southwestern Uganda. Midwifery. 2019;78:16–24. https://doi.org/10.1016/j.midw.2019.07.005.

49. Agustina R, Dartanto T, Sitompul R, Susiloretni KA, Suparmi, Achadi EL, et al. Universal health coverage in Indonesia: concept, progress, and challenges. Lancet. 2019;393(10166):75–102. https://doi.org/10.1016/S0140-6736(18)31647-7.

50. Johar M, Soewondo P, Pujisubekti R, Satrio HK, Adji A. Inequality in access to health care, health insurance and the role of supply factors. Soc Sci Med. 2018;213:134–45. https://doi.org/10.1016/j.socscimed.2018.07.044.

51. Tirgil, A.a B, Gurol-Urganci I, Atun R. Early experience of universal health coverage in Turkey on access to health services for the poor: regression kink design analysis. J Glob Health. 2018;8(2):020412. https://doi.org/10.7189/jogh.08.020412.

52. Miraldo M, Propper C, Williams RI. The impact of publicly subsidised health insurance on access, behavioural risk factors and disease management. Soc Sci Med. 2018;217:135–51. https://doi.org/10.1016/j.socscimed.2018.09.028.

53. Müllerschön J, Koschollek C, Santos-Hövener C, Kuehne A, Müller-Nordhorn J, Bremer V. Impact of health insurance status among migrants from sub-Saharan Africa on access to health care and HIV testing in Germany: A participatory cross-sectional survey 11 Medical and Health Sciences 1117 Public Health and Health Services 11 Medical and Healt. BMC Int Health Hum Rights. 2019;19(1). https://doi.org/10.1186/s12914-019-0189-3.

54. El-Sayed AM, Vail D, Kruk ME. Ineffective insurance in lower and middle income countries is an obstacle to universal health coverage. J Glob Health. 2018;8(2). https://doi.org/10.7189/jogh.08.020402.

55. Chiang C-L, Chen P-C, Huang L-Y, Kuo P-H, Tung Y-C, Liu C-C, et al. Impact of universal health coverage on urban-rural inequity in psychiatric service utilisation for patients with first admission for psychosis: A 10-year nationwide population-based study in Taiwan. BMJ Open. 2016;6(3). https://doi.org/10.1136/bmjopen-2015-010802.

